# Ploidy effects on the relationship between floral phenotype, reproductive investment and fitness exhibited by an autogamous species complex

**DOI:** 10.1101/2022.12.14.520478

**Authors:** Ana García-Muñoz, Camilo Ferrón, Celia Vaca-Benito, João Loureiro, Sílvia Castro, A. Jesús Muñoz-Pajares, Mohamed Abdelaziz

**Affiliations:** Department of Genetics, University of Granada, Granada, Spain; Área de Biodiversidad y Conservación, Departamento de Biología y Geología, Física y Química Inorgánica, Universidad Rey Juan Carlos, Móstoles, Spain; Centre for Functional Ecology, Department of Life Sciences, University of Coimbra, Coimbra, Portugal; Research Unit Modeling Nature, Universidad de Granada, E-18071 Granada, Spain

**Keywords:** *Erysimum*, herkogamy, natural selection, plant phenotype, pollen-ovule ratio, self-pollination, stamen exertion

## Abstract

**Premise:** The relationships between reproductive investment, phenotype and fitness have been broadly studied in cross-pollinated plants in contrast to selfing species, which are considered less interesting in this area because they are supposed to be a dead-end in any evolutionary pathway. Still, selfing plants are unique systems to study these questions since the position of reproductive structures and traits related to flower size play an important role in female and male pollination success.

**Methods:** *Erysimum incanum s*.*l*. is a selfing species complex exhibiting three levels of ploidy: diploids, tetraploids and hexaploids. This species complex shows traits typically associated with the selfing syndrome. Here, we used 1609 plants belonging to these three ploidies to characterize floral phenotype and spatial configuration of reproductive structures, reproductive investment (pollen and ovules production) and plant fitness. Then, we explored the relationship between all these variables using structural equation modelling across ploidy levels.

**Key Results:** An increase in ploidy level leads to bigger flowers with more exerted stamens and a greater amount of pollen and ovules. In addition, hexaploid plants exhibit higher absolute values for herkogamy which is positively correlated with fitness. Phenotypic traits and pollen production are indirectly selected by the relationship among ovules and fitness, maintained across ploidies.

**Conclusions:** Changes in floral phenotypes, reproductive investment and fitness with the ploidy level suggests that genome duplication can be a driver for the reproductive strategy transitions by modifying the investment in pollen and ovules and linking them with plant phenotype and fitness.

## INTRODUCTION

The importance of male and female reproductive investment (i.e., pollen and ovules, respectively) on plant reproduction have been broadly explored in flowering plants (Knight et al., 2005; Morales and Traveset, 2008; Breed et al., 2012; Wong & Frank, 2013). However, most studies have focused mainly on outcrossing plants whose fertilization depends on different pollination mechanisms, leading to a broader understanding of biotic and abiotic interactions in these species. In contrast, selfing plant species have been overlooked in this ecological area, even when a high frequency of self-pollinated hermaphroditic plants occurs in the wild (Jarne and Charlesworth, 1993).

Reproductive strategies based on self-pollination facilitate pollen grains to reach the stigma within the same flower, ensuring fertilization (Lloyd, 1992; Ashman et al., 2004; Eckert et al., 2006). Female reproductive investment is the limiting factor for reproductive success in selfing plants. This causes the reduction of resource allocation to pollen compared to ovules (Lloyd, 1987; Michalski and Durka, 2009). A trade-off between pollen and ovules has been identified, and the P/O ratio was developed as an index to characterize different mating systems (Cruden, 1977). High values of pollen/ovules ratio are expected in outcrossing plant species, where male reproductive investment is a significant component of the plant’s total fitness. Conversely, selfing plant species are expected to show lower pollen/ovules ratio as pollen is only used to fertilize the ovules in their own flower. The pollen/ovule ratio (hereon, P/O ratio) has been broadly used to explore plant mating system evolution and was positively correlated to floral traits driving the evolution of mating systems, such as corolla size or relative spatial position of sexual organs, such as herkogamy (Galloni et al., 2007).

Flower morphology and size were demonstrated long ago to be traits strongly selected in plants as they control the frequency and efficiency of plant reproduction in different taxa (Herrera, 1993; Caruso et al., 2003; 2019), including species in the genus *Erysimum* (Gómez et al., 2006). Morphology and size of flowers can also have a significant effect on the mating system by their relationship with the relative spatial position of sexual organs (i.e., herkogamy; Lloyd and Schoen 1992; Larrinaga et al. 2009; Herlihy and Eckert 2004). Positive herkogamy avoids self-pollination and exposes the stigma surface to pollinators. However, self-pollination is facilitated when herkogamy is negative (reverse herkogamy) or near zero (Larrinaga et al., 2009; Brys et al., 2011). Experimental studies showed how selfing plants often exhibit a lower degree of herkogamy and lower pollen/ovule ratio values because there is a high possibility of own pollen grains reaching the stigma (Johnston et al., 2009; Sicard and Lenhard, 2011). Apart from herkogamy, the stamen exertion can also affect the mating system, which enhances the pollen export by exposing the anthers outside the flower (Barret, 2003, Medano et al., 2005). Because of their relationship with the mating system, evolution in plant traits related to floral morphology, floral size and spatial position between sexual organs are mostly connected to plant fitness mediated by male and female reproductive investment (Cruden, 2000). Thus, the balance between pollen production and ovule development per flower is expected to modify the adaptive trajectories in different populations by constraining the selective scenarios

Another level of complexity in plants is the one produced by whole genome duplications. It is known that about 70% of flowering plants show some degree of polyploidy (Wood et al., 2009). Indeed, polyploidization events play an important role in the evolutionary history of angiosperms (Soltis and Soltis, 1999; Otto and Whitton, 2000; Soltis et al.; 2004; Van der Peer et al.; 2009, Leebens-Mack et al., 2019). Furthermore, ploidy can alter floral phenotypes in different dimensions (Jürgens et al., 2002) and several studies have demonstrated its influence on flower size and position of sexual organs, which, in turn, might affect pollinator preferences (te Beest et al., 2012; Moghe, 2014). Surprisingly, studies often pay poor attention to the effect of ploidy variation on the reproductive output of selfing species, despite it is widely accepted that ploidy level is able to alter floral phenotype in many dimensions (Jürgens et al., 2002).

*Erysimum incanum* is a self-compatible species complex showing small flowers and sharing a similar life form: monocarpic annual plants. The species in the complex differ in their ploidy level (including diploids, tetraploids and hexaploids; Galland, 1988; Luque and Lifante, 1991; Favarger et al., 1979), but they exhibit identical self-pollination mechanisms (Abdelaziz et al., 2019). Due to its self-reproductive strategy, *Erysimum incanum* is expected to increase the allocation of resources to female function compared to male function in its flowers (Charnov, 1982). Because of its short and monocarpic life history, it is possible to measure the entire life production of pollen, ovules and seed output as fitness estimates. Using this singular study system due to its reproductive strategy and its multiploidy condition, the main aims of this study were: (1) to quantify the selective pressures shaping the adaptive trajectories in populations with different ploidy levels; (2) to evaluate the effect of ploidy on the floral patterns resulting from the selective scenarios; (3) to explore the role of pollen/ratio ratio in constraining the total selection regimes occurring through the variation exhibited by the species complex; and (4) evaluate the effects of polyploidization to decouple variation among traits in *E. incanum* species complex. Findings in ecological variation between populations with different ploidy levels might shed light on the role of genetic mechanisms as drivers of the evolutionary trajectory followed by selfing species that generally are considered with a reduced ability to evolve.

## MATERIALS AND METHODS

### Study system

Genus *Erysimum* L. is one of the most diverse in the Brassicaceae family, and their species can be found in Eurasia, North and Central America and North Africa (Al-Shehbaz et al., 2006). Recently, mechanisms promoting local adaptation and hybridization between lineages were identified for different species in the genus (Gómez et al., 2009; Abdelaziz, 2013). *Erysimum incanum* can be considered a species complex, including annual and monocarpic species and subspecies inhabiting the Western Mediterranean basin (Nieto-Feliner, 1993; Abdelaziz et al., 2019). The complex harbours three ploidy levels: diploids (2*n* = 2*x* = 16 chromosomes), tetraploids (2*n* = 4*x* = 32) and hexaplois (2*n* = 6*x* = 48) (Nieto-Feliner, 1993; Abdelaziz et al., in prep.). Diploids of *E. incanum* present a vicariant distribution in the Rif and the Pyrenees mountains and tetraploids present a similar distribution in southwest Iberian Peninsula and the Middle Atlas Mountains (Fennane et al., 1999). In contrast, hexaploid *E. incanum* plants were only found in the most southern ranges in Morocco (High Atlas and Antiatlas; Abdelaziz et al., in prep.) (Fig. 1).

**Figure 1.**
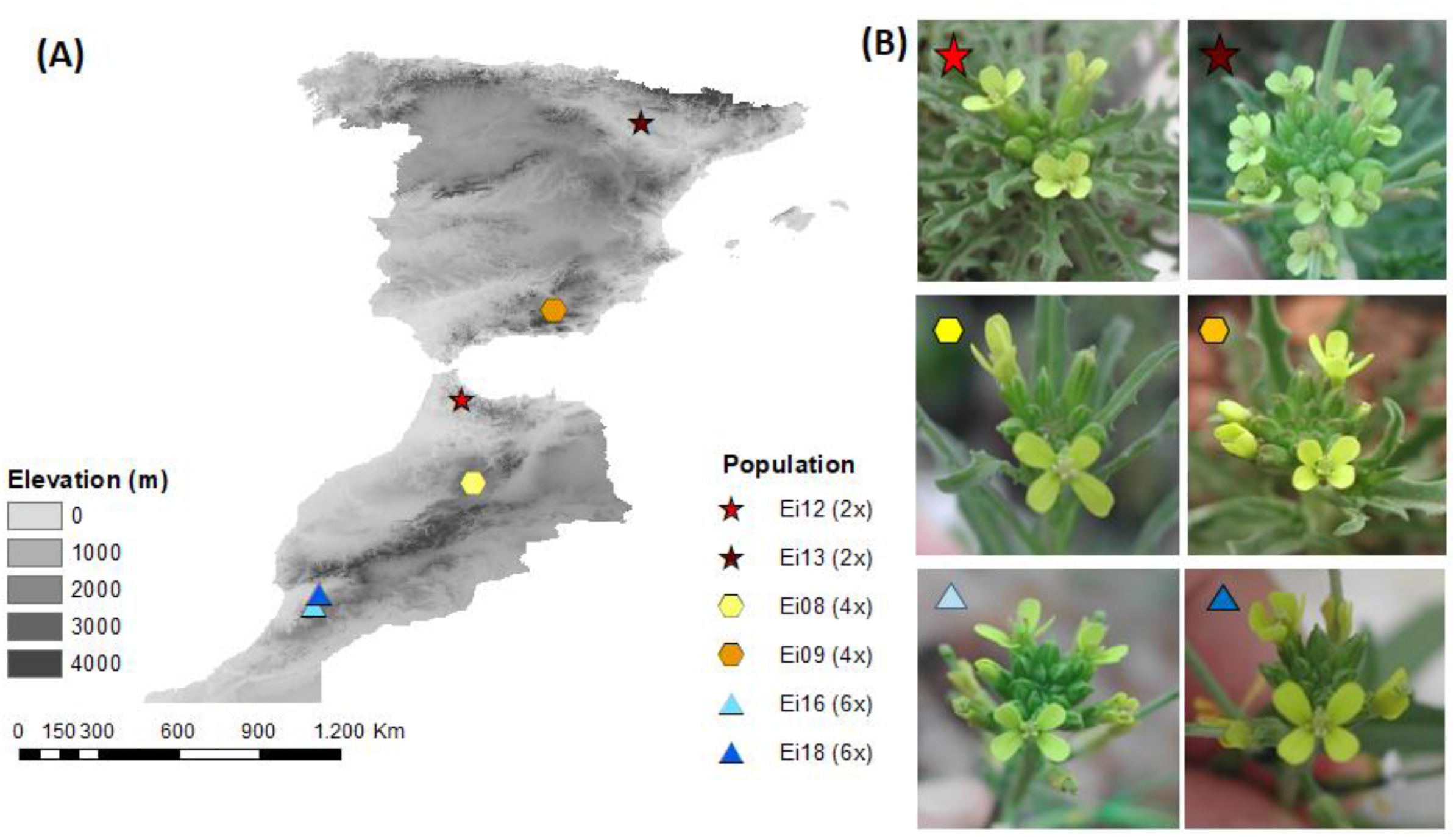
(A) Map of geographic locations of the six studied populations in Morocco and Spain from *Erysimum incanum*. (B) Variation in flower shape and size among different populations and levels of ploidy.

The predominant reproductive strategy of the *E. incanum* species complex is autogamy. Indeed, it shows small, hermaphroditic and self-compatible flowers, where we have recently described the *anther rubbing* mechanism (Abdelaziz et al., 2019) that promotes these plants’ prior selfing.

### Experimental design and phenotypic trait measurements

Seeds collected in two natural populations per ploidy (2*x*: Ei12 and Ei13; 4*x*: Ei08 and Ei09; 6*x*: Ei16 and Ei18; Fig. 1, Table 1) were sowed for three consecutive generations in greenhouse conditions at the University of Granada grow facilities to remove any local effect. A total of 640 seeds per population were sowed in groups of four seeds in square plastic pots (11×11×11 cm^3^) filled with Gramoflor™ potting soil mixture.

**Table 1.** Genome size estimations of each population. The number of plants phenotyped by population and ploidy, the number of plant samples used for genome size analysis, the genome size per population and ploidy (mean ± SE values are shown) and the geographic origin of each studied population are also given.

| Ploidy level | Population | N phenotype | N genome size | Genome size | Geographical origin |
| --- | --- | --- | --- | --- | --- |
| **2x** | Ei 12 | 283 | 21 | 0.35 ± 0.01 | Morocco |
|  | Ei 13 | 114 | 20 | 0.34 ± 0.01 | Spain |
| **4x** | Ei 08 | 513 | 25 | 0.78 ± 0.10 | Morocco |
|  | Ei 09 | 248 | 21 | 0.78 ± 0.03 | Spain |
| **6x** | Ei 16 | 230 | 21 | 1.10 ± 0.03 | Morocco |
|  | Ei 18 | 221 | 20 | 1.13 ± 0.03 | Morocco |

During the blooming period of the plants, we measured a series of floral phenotypic traits on one flower per plant using a digital caliper. Three of these traits were related to floral size, namely petal length (distance between the edge of the petal and the point where the petal starts to curve, shaping the corolla), corolla diameter (distance between the edge of the petal and the diametrically opposite one, in the flower) and corolla tube length (distance between the basis of the sepals and the corolla aperture point). In addition, we also measured the long stamen length (distance between the long filament insertion point and the anther) and the style height (distance between the style basis and the stigma) to estimate the herkogamy value (difference between the style height and the long stamen length) and the stamen exertion (difference between the long stamen length and the corolla tube length). Herkogamy indicates the relative easiness for self-pollination to occur, while stamen exertion denotes how much the anthers (and so the pollen grains) are exposed to flower visitors.

### Genome size measurements

We used young leaf tissue to estimate the genome size and infer the ploidy level using flow cytometry (FCM). We carried out the genome size analyses as described by Muñoz-Pajares et al. (2018). At least 20 individuals were analyzed per population (Table 1). As the ploidy level was homogeneous in each population studied, we assumed that each population was mainly composed of a single cytotype.

### Reproductive investment measurements

As we measured floral traits, half of the stamens were collected (i.e., two long stamens and one short stamen) and conserved in alcohol 70% to estimate male reproductive investment. Next, we counted the number of pollen grains per flower (Pollen) using the Multisizer Coulter Counter 3™ particle counter provided by the morphometric lab of the Centro de Investigación, Tecnología e Innovación (CITIUS) at the University of Seville (Spain).

Once the plant was dried, we collected four matured fruits of each individual to quantify the number of viable and aborted seeds and the number of unfertilized ovules per fruit. The sum of these three measurements is the total number of produced ovules per flower, i.e., the female reproductive investment (Ovules). The pollen-ovule ratio (hereon, P/O ratio) was calculated as the fraction of pollen grains produced per flower divided by the number of ovules produced per plant.

### Fitness estimation

To estimate fitness, we counted the number of fruits per plant and multiplied it by the mean number of seeds estimated using four random fruits from the same plant. This way, we obtained the total seed production per individual plant used in the experiment. This value is an unbiased estimate of the individual fitness due to our plants’ monocarpic character, which completes their life cycle in only 10-12 weeks.

Overall, we considered eight variables grouped into four categories: floral size (petal length, corolla diameter and corolla tube length), the spatial position of reproductive structures (stamen exertion, herkogamy), reproductive investment (pollen, ovules) and fitness. We measured them in a total of 1609 plants (397 diploid, 761 tetraploid and 451 hexaploid plants, according to the ploidy level of the population of origin).

### Statistical analysis

#### Phenotypic traits selection

We used structural equation modelling (SEM) because we are interested in dealing with relationships among multiple variables and test both direct and indirect effects of selection on traits, This statistical technique is a powerful multivariate analysis tool which has gained importance in ecological research in the last two decades (Fan et al., 2016). In addition, SEM incorporates path analysis techniques which are able to identify when a variable effect could be mediated by another different variable. We used SEM to estimate the selection of traits related to floral size, the relative position of reproductive structures, and male and female reproductive investment. Pathway analysis allows us to evaluate the direct and indirect effects of a series of traits that act simultaneously on fitness.

The first stage of SEM is to define the saturated model that will allow us to test the different hypotheses. Based on a priori knowledge from previous studies in related species and in our study system (Gómez et al., 2009; Abdelaziz et al., 2019), we formulated our hypothesis by building the saturated model, as shown in Figure 2. In our model, fitness is a dependent variable directly connected with the rest of the variables in the model. Because reproductive structures, such as stamens and style length, are morphologically dependent on floral size, we included the effect of floral traits on herkogamy and stamen exertion. Flower size traits and the relative position of reproductive organs can promote or avoid self-pollination, which is why we connected them to fitness. All phenotypic traits were also linked with reproductive investment variables to test the influence of floral size and relative position of reproductive structures on pollen and ovules production. Lastly, pollen and ovules were included as having a direct effect on fitness.

**Figure 2.**
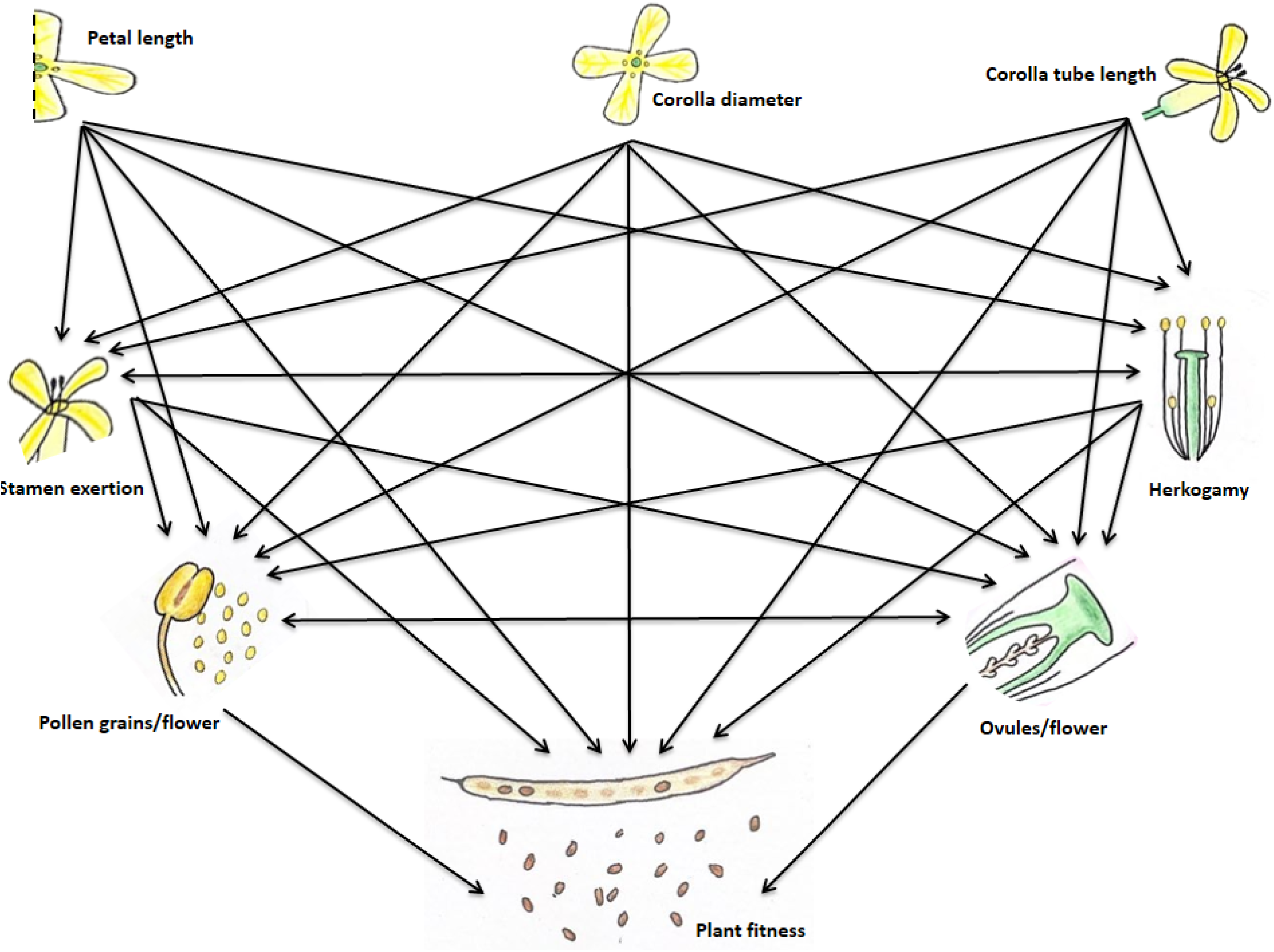
Saturated model hypothesizing the potential relationships among flower traits (petal length, corolla diameter and corolla tube length), relative spatial positions of the sexual organs (stamen exertion and herkogamy), male and female reproductive investment (pollen and ovules) and plant fitness.

Moreover, we considered two potential pairs of covariances. First, we included a covariance between pollen and ovule production, based on the trade-off between male and female sexual allocation (broadly described in flowering plants e.g. Charlesworth and Charlesworth, 1981; Lloyd, 1984; Charnov, 1987), and on the pollen/ovule ratio hypothesis, which suggests that autogamous plant species reallocate resources to increase the production of ovules in detriment of pollen grain production (Cruden, 1977). The second covariance included in our model links herkogamy and stamen exertion. We expect a negative covariance between them due to morphological coherence, i.e., flowers with more exerted stamens would exhibit lower or negative values of herkogamy (reverse herkogamy), unless the stigma is even more exerted than stamens.

The degree of fit between the observed data and the value of the hypothetical model is given by a goodness-of-fit χ^2^. Non-significant χ^2^ indicates we could accept the hypothetical proposed model. To assess the model’s fit to the data, we also considered other indices, such as CFI, which should be near 1, or SRMR, which should be near 0 for a good fit of the model (Smith and McMillan, 2011). To analyse the effect of ploidy variation on trait selection, we fitted the same model to each population and ploidy level separately and compared the resulting different pathways.

#### Trait effects on fitness

Using the relationships of the traits with fitness in the SEM analysis, we calculated the direct, indirect and net values of natural selection acting on each measured phenotypic and reproductive trait. Direct effects result from the unmediated relationship of the traits with fitness. Indirect effects were estimated as the product of different components of a path connecting any trait to fitness mediated by other traits. When multiple independent paths existed for a single trait, they were summed. Finally, we added direct and indirect effects on traits’ net value of natural selection.

#### Effect of P/O ratio on selection regimes

We evaluated the effect of the pollen-ovule ratio on the net selection experienced by each of the phenotypic and reproductive traits, considering the net natural selection regimes experienced by the traits. For this, we performed a regression of the net fitness effect per population on the pollen/ovule ratio.

#### Effect of ploidy level on phenotypic and reproductive investment values

We performed an analysis of variance (ANOVA) followed by Tukey’s HSD test to evaluate possible differences in phenotypic traits, reproductive investment and P/O ratio between the three levels of ploidy exhibited by *E. incanum* species complex. In addition, we assessed for normality assumptions before all statistical analyses and performed transformations (log) of considered variables when necessary.

#### Phenotypic correlation among traits across ploidies

Phenotypic correlations between each pair of *E. incanum* phenotypic traits for each studied ploidies were calculated using Pearson correlation. In addition, we calculate the correlation pooling the plants by ploidy. All the statistical analyses described in this section were conducted in R version 4.0.3, using the package *lavaan* version 0.6-12 for the structural equation modelling, and *stats* for the rest of analyses.

## RESULTS

### Genome size measurements

Flow cytometry analyses showed that the studied populations are homogeneous in their genome size. According to our previous knowledge of genome size variation in the *Erysimum* genus (Nieto-Feliner et al., 1993; Fennane et al., 1999) and knowing the three ploidy levels previously described for the complex, it was possible to assign each individual and population to a specific ploidy level (Table 1).

### Variation in phenotypic correlation among traits

The structural equation modelling comparing populations from each ploidy level showed heterogeneity in the relationships established among traits. However, some patterns are maintained across populations (Fig. 3). We found a significant negative relationship between corolla tube length and stamen exertion in every population, probably because of anatomic coherence. Similarly, we found a negative covariance between stamen exertion and herkogamy across most populations from each ploidy. Since herkogamy exhibits negative values (i.e., it is the difference between the style height and the long stamen length) in this species complex, we must keep in mind that a lower degree of herkogamy translates into a greater separation of male and female structures and a greater exertion of stamens above the corolla.

**Figure 3.**
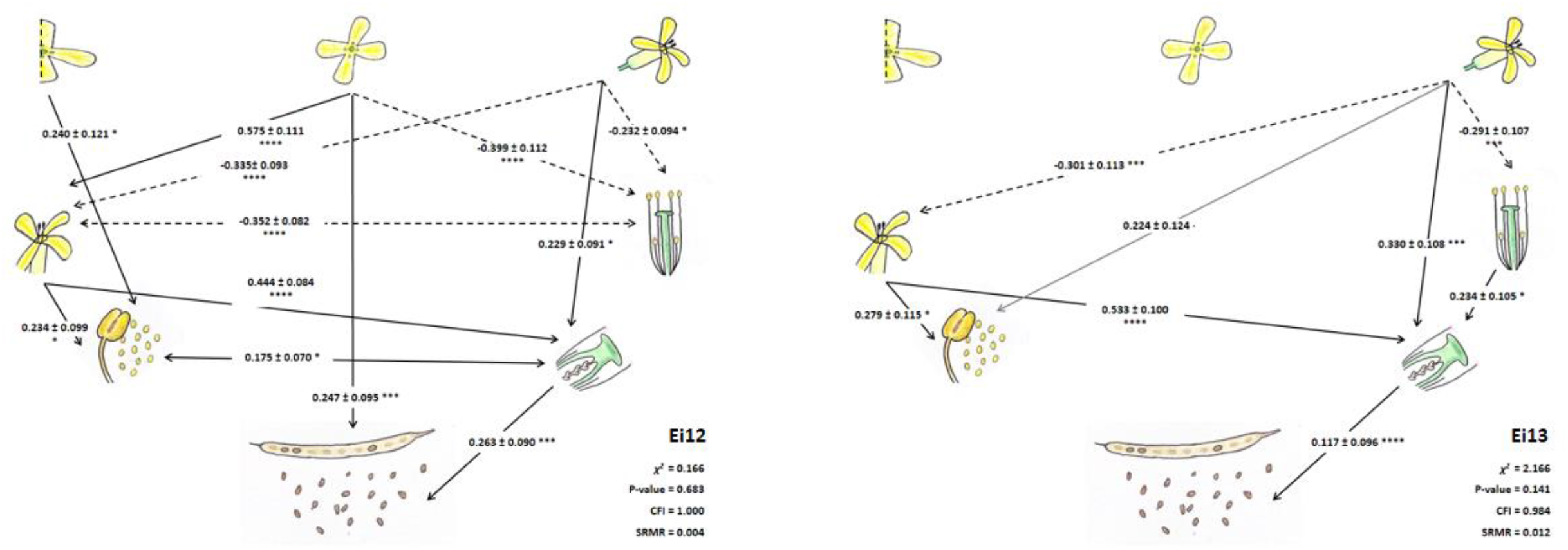
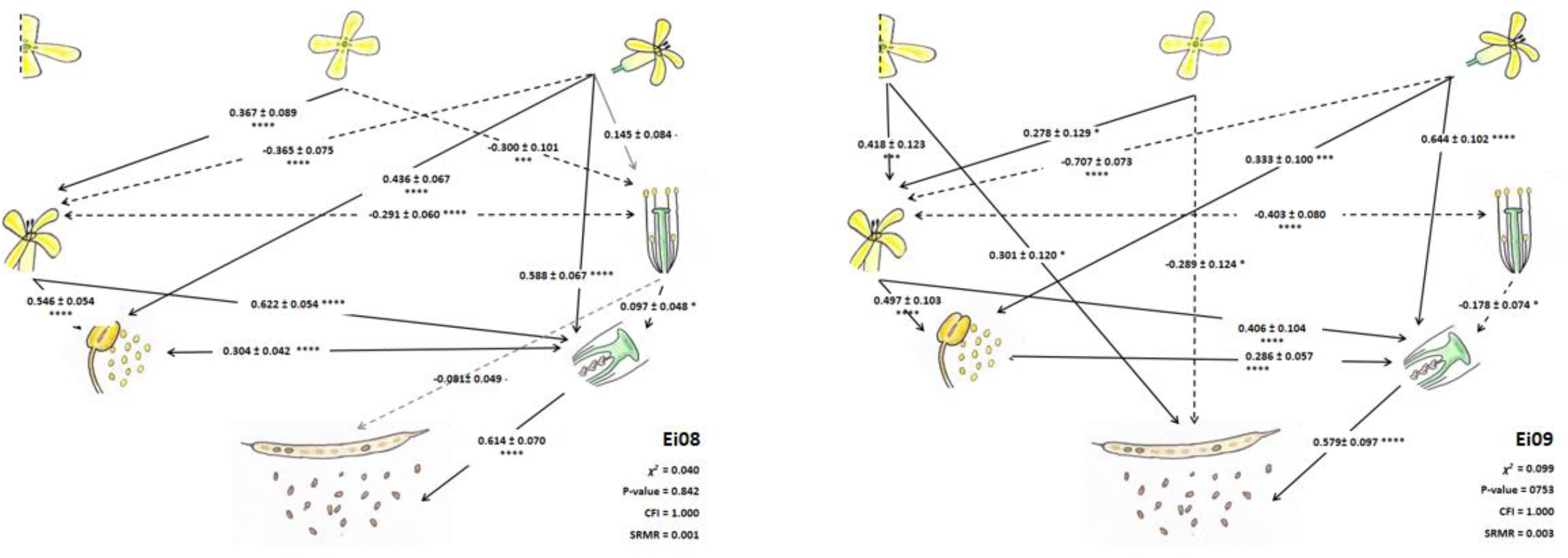
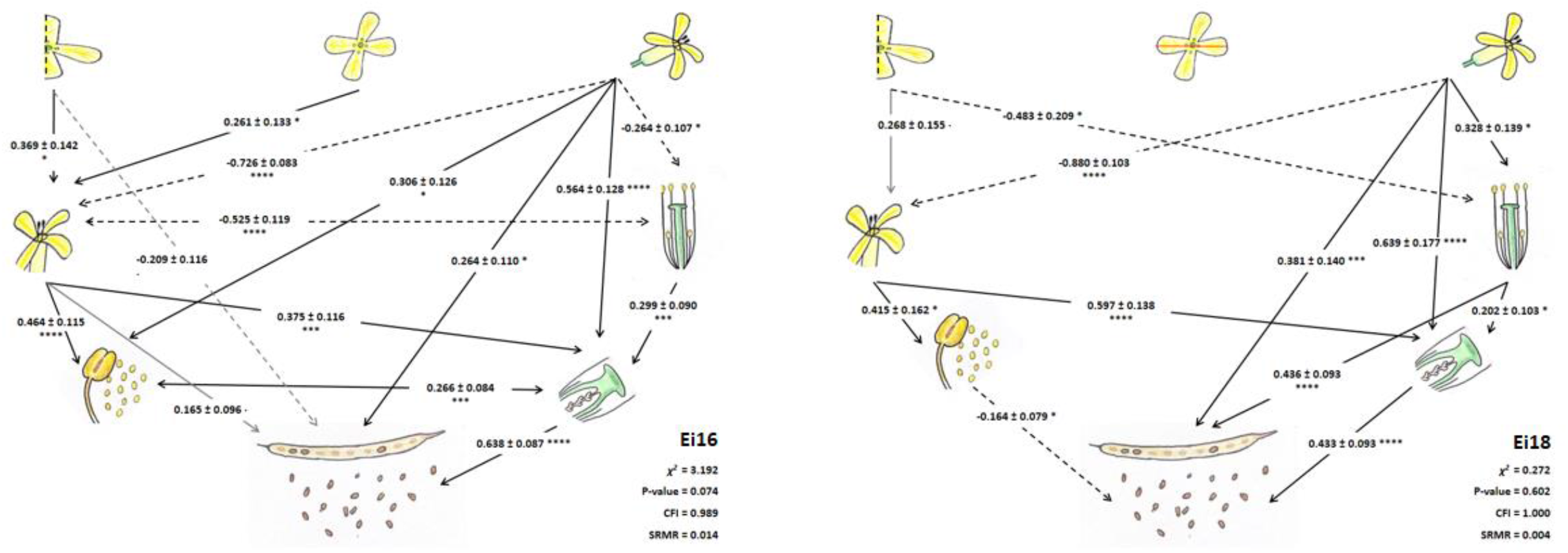
Path diagrams describing the effects of floral phenotypic traits (*petal length, corolla diameter, corolla tube length*), the spatial position of reproductive structures (*stamen exertion* and *herkogamy*) and reproductive investment (pollen and ovules) on fitness for each ploidy level. Solid and dashed arrows indicate positive and negative relationships, respectively. For visual simplicity, only significant relationships are represented, accompanied by the values of path coefficients (mean ± SE).

Apart from stamen exertion, we found that phenotypic traits related to flower size (petal length, corolla diameter and corolla tube length) are commonly negatively connected with herkogamy in all populations except for Ei18, where the relationship is positive. In addition, corolla tube length is positively linked to female reproductive investment (ovules) across all populations. For the male reproductive investment (pollen), we only found a significant link with a floral trait for a diploid population (Ei12) and the tetraploid populations (Fig. 3A and 3B).

Stamen exertion is linked to male and female reproductive investment across all populations, so flowers with more exerted anthers might produce more pollen and ovules. On the other hand, herkogamy is only connected with female reproductive investment. This means that flowers exhibiting less distance between male and female organs could produce more ovules (Fig. 3).

In general, we found similar relationships patterns among traits across populations although the number of significant connections changes. For instance, the lowest number of relationships among traits is shown by the diploid population Ei13 and the number of related traits increases with ploidy level, being maximum for the hexaploid population Ei16.

### Variation in selection on phenotypic and reproductive traits

Patterns of trait selection across populations indicate that ovule amount is the only trait directly linked with fitness in all the studied populations (Fig. 3). Due to this ovule-fitness relationship, traits linked with the female reproductive investment are indirectly selected. Male reproductive investment is directly (and negatively) selected in hexaploid Ei18 only (Fig. 3C). Nevertheless, we found indirect selection acting on pollen mediated by its covariance with ovule amount.

Corolla tube length is under indirect selection in all populations because it is the trait with more established relationships with other traits directly linked with fitness (e.g., the ovules). The other flower size traits are indirectly selected for at least one population from each ploidy level (Fig. 3). Herkogamy is directly selected, while stamen exertion is selected indirectly though ovules and by its covariance with herkogamy (Fig. 3). The estimated parameters and their standard errors resulting from the path analyses per population and ploidy are given in Table S1. The mean values of the direct, indirect and net effect of each studied attribute on fitness are shown in Table S2.

### Effect of P/O ratio on selection regimes

We found a significant relationship between the P/O ratio and selective regimes on herkogamy across populations (Fig. 4). The proportion of variance of selective regimes explained by the P/O ratio (R^2^ = 0.73) was significant (*P =* 0.03) even when the number of populations included in the analysis was low. In populations where the P/O ratio was low, the approximation between stamens and stigma is selected. While in populations where the P/O ratio is higher, the separation between anther and stigma is positively selected (Fig. 4 and Table 2).

**Table 2.** Total selective effects on traits resulting from the sum of direct effects and the product from mediated effects on fitness according to SEM for each population. Significant values are shown in bold, while the other values result from the product of significant and marginally significant effects.

| Traits | 2n |  | 4n |  | 6n |  |
| --- | --- | --- | --- | --- | --- | --- |
|  | Ei12 | Ei13 | Ei08 | Ei09 | Ei16 | Ei18 |
| Petal length | **0.011** |  |  | **0.451** | -0.129 | -0.253 |
| Corolla diameter | **0.338** |  | 0.272 | **-0.190** | **0.057** |  |
| Corolla tube length | **0.028** | **0.012** | **0.196** | **0.174** | **0.489** | **0.386** |
| Stamen exertion | **0.128** | **0.062** | **0.456** | **0.359** | **0.382** | **0.190** |
| Herkogamy | **-0.043** | **0.027** | -0.133 | **-0.231** | **0.065** | **0.523** |
| Pollen | **0.046** |  | **0.187** | **0.166** | **0.170** | **-0.164** |
| Ovules | **0.263** | **0.117** | **0.614** | **0.579** | **0.638** | **0.433** |

**Figure 4.**
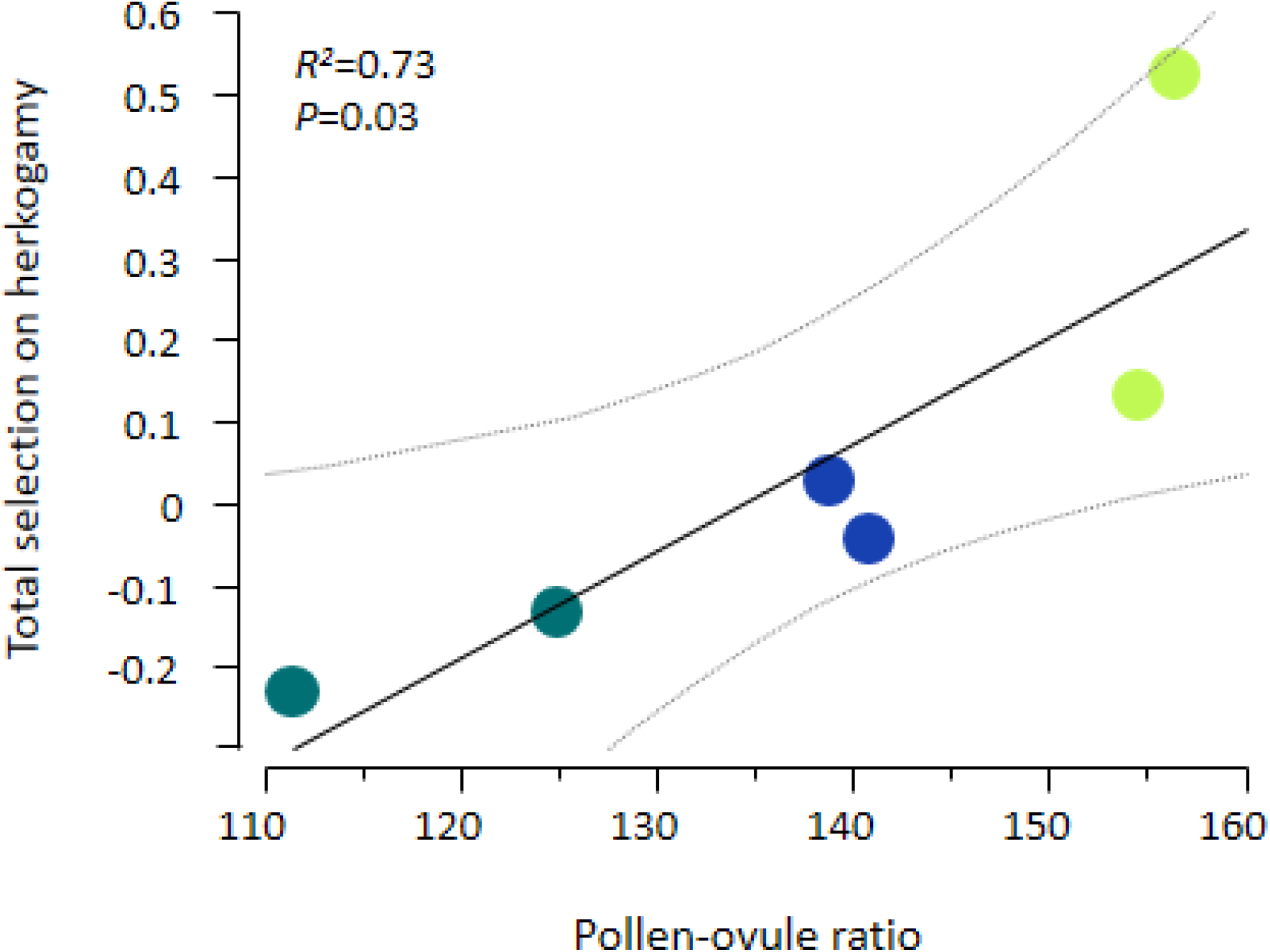
Effect of P/O ratio on selective strength. Response curves show the P/O ratio effect on the total selection on herkogamy. Each circle represents a single population: green circles refer to hexaploid populations, blue circles to diploid populations and ocean circles to tetraploid populations. Grey lines represent 95% confidence intervals.

### Effect of ploidy level on phenotypic and reproductive traits

We found significant among-ploidy differences in all traits related to floral size (petal length, corolla diameter, corolla tube length) separately (Fig. 5). Plants with higher ploidy showed bigger corollas, and this pattern is maintained for the corolla tube length.

**Figure 5.**
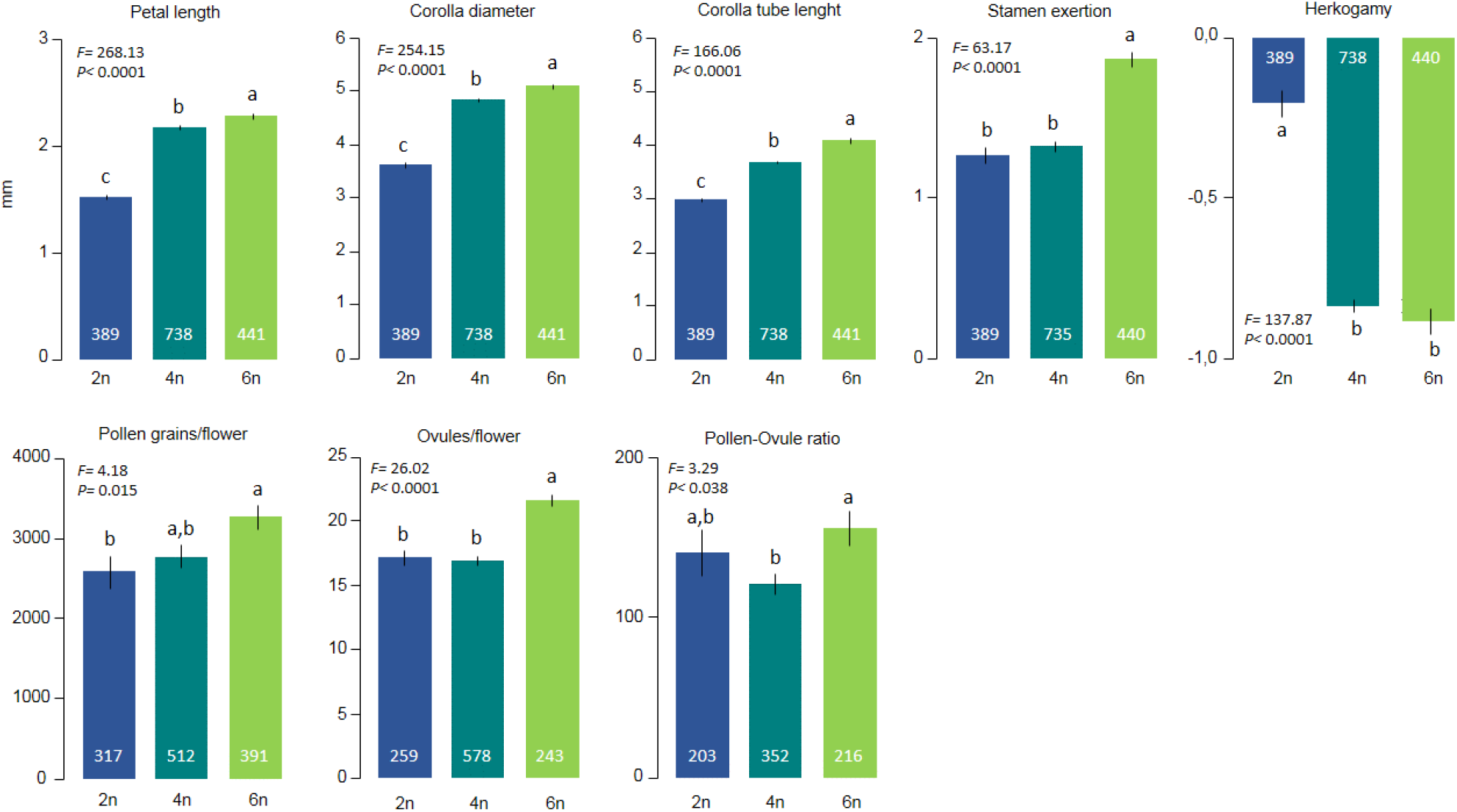
Mean and SE values for petal length, corolla diameter, corolla tube length, stamen exertion, herkogamy, pollen and ovules amount per flower and pollen/ovule ratio for each ploidy level. F ratios refer to one-way ANOVA, and numbers inside the bars indicate the number of individuals used for each measured trait. Letters refer to these groups that are statistically significant according to a Tukey HSD comparison.

Flowers of hexaploid plants have significantly more exerted stamens above the corolla compared with diploid and tetraploid plants. We found significant differences among diploid and polyploid plants. Diploid plants exhibit reduced values of herkogamy compared to tetraploid and hexaploid plants, which present more negative values of herkogamy (Fig. 5).

Regarding reproductive investment, we found significant differences among ploidy levels but not the same pattern for pollen and ovules. Hexaploid plants produce a significantly higher amount of pollen and diploids showed the lowest value, with the tetraploids presenting an intermediate value (Fig. 5). However, hexaploids exhibited significantly higher values of ovules per flower diploids and tetraploids that showed no significant differences for this trait (Fig. 5). When we evaluated the P/O ratio, hexaploids presented the highest values. Still, diploids showed intermediate values and tetraploids the lowest ones (Fig. 5).

### Phenotypic correlation among traits across ploidies

We found significant differences between the phenotypic correlations among traits across ploidy levels. Diploids exhibited significant correlations between every pair of measured traits. However, the number of significant correlations decreased in tetraploids and dropped even more in number when we analyzed hexaploids (Table 3). There was a significant positive phenotypic correlation among all the attributes except for herkogamy, which negatively correlated with all the other traits. In tetraploids, most correlations were significant and positive; however, any correlation with herkogamy and the correlation between corolla tube length and exertion were negative. The correlations between corolla diameter-exertion and corolla tube length-herkogamy were non-significant. Finally, we found significant positive correlations in hexaploids between petal length, corolla diameter and corolla tube length. These floral attributes also showed significant positive correlations with ovules. Corolla tube length and herkogamy, exertion and pollen grains, and pollen grains and ovules were also significant and positively correlated. There were significant negative correlations of exertion with petal length, corolla tube length and herkogamy, and between herkogamy and pollen production. The rest of phenotypic correlations were non-significant in hexaploid plants (Table 3).

**Table 3.** Phenotypic correlations among *Erysimum incanum* phenotypic traits for each ploidy level. Positive correlations are highlighted in blue, while negative correlations are highlighted in red. The sample size is shown by ploidy. Significance levels after Bonferroni corrections: **P < 0.01; ***P < 0.001; ****P < 0.0001, are also provided.

| 2n (397) | Corolla diameter | Corolla tube length | Exertion | Herkogamy | Pollen | Ovules |
| --- | --- | --- | --- | --- | --- | --- |
| Petal length | **0.7908\*\*\*\*** | **0.6473\*\*\*\*** | **0.3244\*\*\*\*** | **-0.3493\*\*\*\*** | **0.2091\*\*\*\*** | **0.4619\*\*\*\*** |
| Corolla diameter |  | **0.6140\*\*\*\*** | **0.4343\*\*\*\*** | **-0.4103\*\*\*\*** | **0.1770\*\*** | **0.5513\*\*\*\*** |
| Corolla tube length |  |  | **0.1965\*\*\*\*** | **-0.3872\*\*\*\*** | **0.1677\*\*** | **0.5558\*\*\*\*** |
| Exertion |  |  |  | **-0.4908\*\*\*\*** | **0.3563\*\*\*\*** | **0.6124\*\*\*\*** |
| Herkogamy |  |  |  |  | **-0.2787\*\*\*\*** | **-0.4195\*\*\*\*** |
| Pollen |  |  |  |  |  | **0.3263\*\*\*\*** |
| 4n (761) | Corolla diameter | Corolla tube length | Exertion | Herkogamy | Pollen | Ovules |
| Petal length | **0.6534\*\*\*\*** | **0.5491\*\*\*\*** | **0.1013\*\*** | **-0.1011\*\*** | **0.2703\*\*\*\*** | **0.3490\*\*\*\*** |
| Corolla diameter |  | **0.5109\*\*\*\*** | 0.0874 | **-0.1023\*\*** | **0.2420\*\*\*\*** | **0.3122\*\*\*\*** |
| Corolla tube length |  |  | **-0.2622\*\*\*\*** | -0.0214 | **0.1480\*\*\*** | **0.4095\*\*\*\*** |
| Exertion |  |  |  | **-0.3482\*\*\*\*** | **0.4420\*\*\*\*** | **0.3418\*\*\*\*** |
| Herkogamy |  |  |  |  | **-0.1906\*\*\*\*** | **-0.1887\*\*\*\*** |
| Pollen |  |  |  |  |  | **0.6691\*\*\*\*** |
| 6n (451) | Corolla diameter | Corolla tube length | Exertion | Herkogamy | Pollen | Ovules |
| Petal length | **0.6956\*\*\*\*** | **0.5492\*\*\*\*** | **-0.1456\*\*** | 0.0132 | 0.0544 | **0.2755\*\*\*\*** |
| Corolla diameter |  | **0.3668\*\*\*\*** | -0.0109 | 0.0417 | -0.0614 | **0.2134\*\*** |
| Corolla tube length |  |  | **-0.5936\*\*\*\*** | **0.1248\*\*** | -0.0509 | **0.2855\*\*\*\*** |
| Exertion |  |  |  | **-0.2226\*\*\*\*** | **0.2622\*\*\*\*** | 0.1174 |
| Herkogamy |  |  |  |  | **-0.1408\*\*** | 0.0627 |
| Pollen |  |  |  |  |  | **0.2861\*\*\*\*** |

## DISCUSSION

The efficacy of natural selection acting on a selfing plant species has been explored at a molecular level (Arunkumar et al., 2015) but received less attention than outcrossing species, mainly when the focus was natural selection acting on phenotypic traits or reproductive investment. By combining the study of relationships among traits, reproductive investment, and their direct and indirect effects on fitness across different ploidy levels, this work shows that changes in the DNA amount significantly affect the configuration of the relationships established between the studied floral traits, modifying the phenotypic integration of these traits across ploidies. In this group of selfing species, the effect of flower traits on reproductive investment and self-pollination plays a vital role in plant fitness, shaping the floral pattern we observe across ploidies. However, the strong selection of traits related to pollen export makes them key attributes driving changes in the evolution of reproductive strategies within a selfing multiploidy species complex.

Path analyses made it possible to detect the direct, indirect and net effects of traits on fitness. This is essential because indirect effects frequently pass undetected, as traits interact with fitness and several selective agents simultaneously, which are not always under our control. Often, traits are mediated by natural selection on other traits, modifying the selective pressure. For example, Gómez et al. (2009), using a SEM approximation on *Erysimum mediohispanicum* populations, described significant effects of plant phenotype by indirect selection on the total selection experienced by the plants. This work demonstrated that this indirect selection played a crucial role in the rise of the geographic mosaic of selection in *E. mediohispanicum* in the Sierra Nevada Mountains.

In our case, we found that traits related to corolla size are scarcely under direct selection but more frequently under indirect selection. Cost production and maintenance of the structure could explain the negative selection acting on some of these traits (Ashman and Schoen, 1994). However, bigger flowers enhance the exertion of the stamens, which has a positive effect on fitness (Gómez et al., 2009; Sahli and Conner, 2011). Evaluating the net effect on fitness, we can observe an existing counterbalance between different selective gradients. Previous studies have already show that opposite selective pressures acting on the same trait are important to maintain phenotypic variation within a population (Siepielski and Benkman, 2010), which is essential for evolution. Additionally, differential responses to selection by the same trait could favour diverse evolutionary trends within a species (Zhou et al., 2020). Consequently, if the selected traits are related to the reproductive strategy, a transition in the mating system might be possible in the *E. incanum* complex.

Polyploidization has the potential to produce changes in flower morphology, size and physiology (Stebbins, 1971; Müntzing, 1936; Levin, 2002; Clo and Kolàr, 2021). Recent works comparing diploid and polyploid populations showed that higher ploidy levels exhibit not only larger flowers but also larger pollen grains (Oliveira et al., 2022). In addition, studies in *Heuchera grossularifolia* (Segraves and Thompson, 1999) demonstrated that tetraploid populations showed larger flowers conducting to a different pollinator assemblage than diploid populations. We also found a tendency for bigger flowers and higher levels of pollen grain production as ploidy increases in *E. incanum*. Although we did not test the existence of changes in pollinator visits, larger flowers associated with higher pollen production could increase floral advertisement and attraction, modifying pollinator interactions. Changes in metabolic routes could accompany these changes in ploidy level and flower size. An interesting example of this has been described in the *Dianthus broteri* species complex, where an increasing floral scent variation was associated with polyploidization (Picazo-Aragonés et al., 2020). In *E. incanum*, it would be interesting to evaluate changes in metabolites related to this increment in flower size and pollen grain production as a potential mechanism to attract flower visitors.

Anther’s length play an important role in the exposure (and subsequent pollen removal) of self-pollen and the reception of outcrossing pollen grains by attracting pollen vectors. In autogamous plants, Rosas-Guerrero et al. (2010) demonstrated a high degree of integration among floral traits that promote self-pollination, such as the style and anthers lengths. In addition, higher values of stamens and pistil length are associated with greater pollen exposure and stronger competition between self-pollen and the incoming outcrossing pollen, as shown by classical studies performed in animal-pollinated plants (Campbell, 1989). Yet, anther’s length does not necessarily relate with flower size, as observed here. According to our results, and contrasting with other autogamous mating systems, in *E. incanum* high values of floral traits, pollen amount and exertion in polyploids could suggest outcrossing reproduction might be gaining importance with increased ploidy level.

Herkogamy is one of the primary mechanisms reducing selfing and driving mating system transitions in plants (Opedal, 2017). *Erysimum incanum* exhibits reverse or negative herkogamy, i.e., the stamens are above the stigma surface to ensure self-pollen deposition in the stigma. However, reverse herkogamy is more accentuated when ploidy increases, with the biggest distances among sexual organs being observed in the hexaploids. Increasing values of herkogamy together with higher anther exertion would promote pollen exportation more easily after pollinator visits, even when an anther-rubbing mechanism occurs in *E. incanum* (Abdelaziz et al., 2019) and self-pollination is always assured.

It is worth noting that reproductive investment indirectly mediates most traits’ selection. This is especially clear when we focus on ovule production, as the most significant selection of traits was exclusively mediated via female function. In addition, we found variability in pollen production associated with different phenotypes and significant differences between ploidy levels. So, to understand the full scope of these changes in sexual effort, we considered focusing on the P/O ratio and its evolutionary consequences. According to the sex allocation theory (Charnov, 1987; Brunet, 1992; Campbell, 2000), we expect a trade-off between male and female reproductive investment. Cruden (1977) described a trade-off mechanism between pollen and ovules production by proposing that low values in the P/O ratio are associated with autogamous mating systems in plants, while high values would be characteristic of outcrossing species. This pattern has been described in *Silene* and *Dianthus* species, where lower levels of P/O ratio were associated with self-compatible species (Jürgens et al. 2002). However, the positive covariance between pollen and ovule production we found in our system means that plants investing more in male reproduction also invest more in female reproduction. Our results demonstrate that a trade-off between male and female reproductive investment does not exist in *Erysimum incanum*. This finding is unexpected for a selfing species.

We also detected a positive effect of the P/O ratio on the natural selection intensity exhibited on herkogamy across the analyzed populations. This result suggests that in populations with a lower P/O ratio, an approximation of the anthers to stigma is favored. Nevertheless, in populations where the P/O ratio is higher, the separation between male and female reproductive organs is favored by selection. This outcome agrees with previous results in other species from the genus *Melochia* (Faife-Cabrera et al., 2018) and Fabaceae species (Galloni et al., 2007), where a positive correlation was described between P/O ratio and herkogamy. Our findings show that in populations where the proximity between male and female reproductive organs is selected, the P/O ratio decreases, probably because self-pollination is more efficient. Conversely, in a population where the separation between sexual organs is strongly selected, the P/O ratio increases, likely to increase the success of outcrossing. These findings support and give a mechanistic explanation to Cruden’s conclusion that the P/O ratio decreases with an efficient degree of self-fertilization (Cruden, 1977).

Studying multiple traits affecting fitness allowed us to have a global view of the different selecting pressures acting on phenotypes. But it also allowed us to explore the possible relationship between the measured traits and their integration into the phenotype. In this sense, the graduate reduction of significant correlations from the diploid to hexaploid plants suggests that the increasing ploidy level significantly affects the covariation between plant traits (i.e., phenotypic integration). Phenotypic integration was pointed out as adaptive when it results from convergent evolution in the values of functional traits to develop a common or independent function in the organism. However, we can also think of phenotypic integration as a constraint for future evolution of each trait’s covariance (Pigliucci, 2003). For example, sets of vegetative or reproductive attributes sharing common functions would strongly covary and hence limit the independent variation of each other. Polyploidy was described as contributing to modify covariation among traits in *Dianthus broteri* species complex (Balao et al., 2011) and *Brassica* allopolyploid species (Baker et al., 2017) and now in *E. incanum* complex as well (results herein). Reduced covariation values between traits may reduce and overcome the constraints imposed by phenotypic integration, allowing the new polyploid species to explore new trait spaces and potentially reach new or higher adaptive peaks unreached by their diploids counterparts.

## CONCLUSIONS

Changes in flower size, spatial position of sexual organs, and reproductive investment seem to be a direct consequence of genome duplications, with hexaploid plants showing the highest values for every studied attribute. No trade-off was found between pollen and ovule production. Still, the number of ovules per flower was the trait under significant selective pressure in every population independently of the ploidy. This evidences the limiting role of ovules as a fitness component for these plants. In addition, ovules also significantly affected adaptive trajectories by their relation with pollen production (P/O ratio), which constrains the selective pressures acting on important traits such as herkogamy. Finally, increasing ploidy values contributed significantly to decoupling variation among traits in *E. incanum* species complex, reducing phenotypic integration as ploidy increases. So, even working with a selfing species complex, our results evidence that the variation promoted by polyploidization is a key factor in the reproductive strategy transitions by modifying the investment in pollen and ovules and linking them with plant fitness and phenotype evolution.

## ACKNOWLEDGMENTS

The authors thank Modesto Berbel, MariPaz Solis, Andrea Martín Salas, Cristobal Bragagnolo, and Mariana Castro for lab assistance and paper discussion at the research group. We also thank Luis Matías and María José Ariza from University of Sevilla for their helpful assistance and management to work in CITIUS. This research has been supported by a grant from the Spanish Ministry of Economy and Competitiveness (CGL2014-59886-JIN), the Organismo Autónomo de Parques Nacionales (Ref: 2415/2017), and the Ministry of Science and Innovation (PID2019-111294GB-I00/SRA/10.13039/501100011033), including FEDER funds. AJM-P was funded by the European Commission under the Marie Sklodowska-Curie Action Cofund 2016 EU agreement 754446 and the UGR Research and Knowledge Transfer—Athenea3i. AG-M was supported by the *OUTevolution* project (PID2019-111294GB-I00/SRA/10.13039/501100011033).

## AUTHOR CONTRIBUTIONS

AGM, CF, AJMP and MA thought and designed the experiments. AGM, CF and CVB conducted the greenhouse experiments. JL and SC conducted the ploidy analyses. AGM, AJMP and MA made the statistical analyses and designed the tables and figures. AGM wrote the first draft of this manuscript and all the rest of authors made significant contribution to the draft. AJMP, SC and MA got the funds to develop this study. AJMP and MA supervised all the study.

## DATA AVAILABILITY STATEMENT

The original contributions presented in the study are included in the article Supplementary Material. Further inquiries can be directed to the corresponding author/s.

