## Supplementary material for "Ploidy effects on the relationship between floral phenotype, reproductive investment and fitness exhibited by an autogamous species complex": Table S1

| 2x | Stamen exertion | Herkogamy | Pollen | Ovules | Fitness |
|---|---|---|---|---|---|
| **Petal length** | -0.019 ± 0.079 | 0.022 ± 0.080 | 0.076 ± 0.081 | **-0.158 ± 0.077*** | 0.072 ± 0.076 |
| **Corolla diameter** | **0.559 ± 0.075 ****** | **-0.311 ± 0.077 ****** | -0.078 ± 0.083 | **0.239 ± 0.079 **** | 0.049 ± 0.079 |
| **Corolla tube length** | **-0.174 ± 0.063 ***** | **-0.180 ± 0.064 **** | 0.064 ± 0.067 | **0.210 ± 0.063 **** | 0.100 ± 0.064 |
| **Stamen exertion** | | **-0.312 ± 0.045 ****** | **0.322 ± 0.058 ****** | **0.331 ± 0.055 ****** | 0.051 ± 0.059 |
| **Herkogamy** | | | **-0.149 ± 0.057 **** | -0.007 ± 0.054 | 0.083 ± 0.054 |
| **Pollen** | | | | -0.036 ± 0.041 | 0.021 ± 0.049 |
| **Ovules** | | | | | **0.429 ± 0.077 ****** |

| 4x | Stamen exertion | Herkogamy | Pollen | Ovules | Fitness |
|---|---|---|---|---|---|
| **Petal length** | **0.277 ± 0.050 ****** | -0.063 ± 0.055 | **-0.106 ± 0.048 *** | -0.037 ± 0.049 | 0.016 ± 0.044 |
| **Corolla diameter** | **0.152 ± 0.049 ***** | -0.088 ± 0.054 | 0.067 ± 0.047 | 0.073 ± 0.048 | 0.050 ± 0.044 |
| **Corolla tube length** | **-0.492 ± 0.044 ****** | 0.072 ± 0.048 | **0.400 ± 0.045 ****** | **0.379 ± 0.046 ****** | **0.086 ± 0.032 *** |
| **Stamen exertion** | | **-0.298 ± 0.037 ****** | **0.467 ± 0.038 ****** | **0.467 ± 0.038 ****** | **0.382 ± 0.039 ****** |
| **Herkogamy** | | | -0.026 ± 0.035 | -0.053 ± 0.036 | **-0.086 ± 0.032 **** |
| **Pollen** | | | | **0.195 ± 0.030 ****** | **-0.089 ± 0.035 *** |
| **Ovules** | | | | | **0.568 ± 0.035 ****** |

| 6x | Stamen exertion | Herkogamy | Pollen | Ovules | Fitness |
|---|---|---|---|---|---|
| **Petal length** | **0.257 ± 0.048 ****** | -0.086 ± 0.052 | -0.043 ± 0.046 | -0.031 ± 0.047 | 0.010 ± 0.042 |
| **Corolla diameter** | **0.176 ± 0.047 ****** | -0.086 ± 0.051 | 0.059 ± 0.044 | 0.069 ± 0.045 | **-0.103 ± 0.041 *** |
| **Corolla tube length** | **-0.509 ± 0.043 ****** | **0.113 ± 0.047 *** | **0.295 ± 0.034 ****** | **0.379 ± 0.046 ****** | **0.131 ± 0.038 ***** |
| **Stamen exertion** | | **-0.305 ± 0.036 ****** | **0.555 ± 0.038 ****** | **0.386 ± 0.039 ****** | -0.053 ± 0.040 |
| **Herkogamy** | | | 0.039 ± 0.034 | -0.028 ± 0.035 | -0.043 ± 0.030 |
| **Pollen** | | | | **0.263 ± 0.030 ****** | **0.116 ± 0.034 ***** |
| **Ovules** | | | | | **0.536 ± 0.036 ****** |

Table S1. Estimate parameters and standard error of direct effects on reproductive structures position (Stamen exertion and Herkogamy) and male and female reproductive investment (Pollen and Ovules, respectively) based on Structural Equation Modelling for each ploidy level. Values in italics refer to covariance estimates. Significant relationships and the significant degree are shown in bold.

*P < 0.05, **P < 0.01, ***P < 0.001, ****P < 0.0001.
