## Supplementary material for "Ploidy effects on the relationship between floral phenotype, reproductive investment and fitness exhibited by an autogamous species complex": Table S2

| Traits | | Ei08 | | | Ei09 | |
|---|---|---|---|---|---|---|
| | **Direct effect** | **Indirect effect** | **Net effect** | **Direct effect** | **Indirect effect** | **Net effect** |
| **Petal length** | | | | 0.301 | 0.150 | 0.451 |
| **Corolla diameter** | | 0.272 | 0.272 | -0.289 | 0.099 | -0.190 |
| **Corolla tube length** | | 0.196 | 0.196 | | 0.174 | 0.174 |
| **Stamen exertion** | | 0.456 | 0.456 | | 0.359 | 0.359 |
| **Herkogamy** | -0.081 | -0.081 | -0.162 | | -0.231 | -0.231 |
| **Pollen** | | 0.187 | 0.187 | | 0.166 | 0.166 |
| **Ovules** | 0.614 | | 0.614 | 0.579 | | 0.579 |

| Traits | | Ei12 | | | Ei13 | |
|---|---|---|---|---|---|---|
| | **Direct effect** | **Indirect effect** | **Net effect** | **Direct effect** | **Indirect effect** | **Net effect** |
| **Petal length** | | 0.011 | 0.011 | | | |
| **Corolla diameter** | 0.247 | 0.091 | 0.338 | | | |
| **Corolla tube length** | | 0.028 | 0.028 | | 0.012 | 0.012 |
| **Stamen exertion** | | 0.128 | 0.128 | | 0.062 | 0.062 |
| **Herkogamy** | | -0.043 | -0.043 | | 0.027 | 0.027 |
| **Pollen** | | 0.046 | 0.046 | | | |
| **Ovules** | 0.263 | | 0.263 | 0.117 | | 0.117 |

| | | Ei16 | | | Ei18 | |
|---|---|---|---|---|---|---|
| **Traits** | **Direct effect** | **Indirect effect** | **Net effect** | **Direct effect** | **Indirect effect** | **Net effect** |
| **Petal length** | | 0.080 | 0.080 | | -0.253 | -0.253 |
| **Corolla diameter** | | 0.057 | 0.057 | 0.381 | 0.281 | 0.662 |
| **Corolla tube length** | 0.264 | 0.225 | 0.489 | | 0.252 | 0.252 |
| **Stamen exertion** | 0.165 | 0.217 | 0.382 | | 0.190 | 0.190 |
| **Herkogamy** | | 0.024 | 0.024 | 0.436 | 0.087 | 0.523 |
| **Pollen** | | 0.170 | 0.170 | -0.164 | | -0.164 |
| **Ovules** | 0.638 | | 0.638 | 0.433 | | 0.433 |

Table S2. Mean values of each trait's direct and indirect and net effect on fitness for each population. Indirect effects are calculated from the mediated effects on fitness product, based on estimates of structural equation modelling. The net effect is the sum of direct and indirect effects.
